# ChatMDV: Democratising Bioinformatics Analysis Using Large Language Models

**DOI:** 10.1101/2025.08.26.671083

**Authors:** Maria Kiourlappou, Peter Todd, Yaxuan Kong, Jayesh Hire, Sibgathullah Furquan Nawab Mohammed, Martin Sergeant, Stefan Zohren, Jim Hughes, Stephen Taylor

## Abstract

**Background:** The rapid advancement in single-cell, spatial omics, imaging, and genomic technologies requires robust analytical and visualisation platforms capable of managing complex biological data. Tools such as Multi-Dimensional Viewer (MDV) offer comprehensive interfaces for data exploration, but still require manual configuration and computational expertise to generate visualisation outputs, limiting accessibility for many users.

**Results:** We present ChatMDV, a natural language interface integrated with MDV that allows users to generate high-quality interactive visualisations through natural language commands. ChatMDV employs a retrieval-augmented generation (RAG) pipeline combined with large language models (LLMs) to translate user queries into reproducible Python code and interactive output. This approach enables exploratory and targeted analysis in diverse biological domains. We demonstrate ChatMDV’s capabilities using three datasets of increasing complexity: the Peripheral Blood Mononuclear Cells 3K (PBMC3K) dataset, the lung cancer atlas dataset hosted at the Human Cell Atlas and the longitudinal TAURUS study single-cell RNA-sequencing (scRNA-seq) dataset.

**Conclusions:** By bridging the gap between natural language processing and bioinformatics visualisation, ChatMDV reduces technical barriers, enhances reproducibility, and supports more inclusive scientific inquiry. Its modular design and adherence to FAIR (Findability, Accessibility, Interoperability, and Reuse) principles make it a scalable and adaptable framework for accelerating biological data analysis.

**Key Points:**

- ChatMDV enables users to create interactive visualisations from biological datasets using natural language.
- The system combines large language models with MDV’s graphical platform to simplify data exploration.
- It supports reproducibility, adaptability, and FAIR data practices, making it suitable for a wide range of users and use cases.

## Background

The increasing scale and complexity of high-throughput biological data, particularly from single-cell, spatial, and imaging omics, has created a critical need for accessible and scalable visualisation tools. Recent advances in imaging and sequencing technologies have allowed researchers to interrogate biological systems with unprecedented resolution.

Although platforms such as Vitessce [1], Napari [2], and Multi-Dimensional Viewer (MDV) [3] have significantly advanced interactive exploration of multimodal data, many of these tools require manual configuration or programming knowledge, creating barriers for researchers without computational training. In particular, Vitessce [1] provides web-based linked views for spatial and singlecell datasets via an intuitive point-and-click interface; however, its customisation and layout definitions require manual editing of JSON configuration files, limiting flexibility for non-technical users. Napari, excels in n-dimensional microscopy data visualisation, offering a python based desktop interface for image segmentation, annotation, and plugin-based analysis. While it is extensible and performant, it is primarily optimised for image data rather than tabular or multimodal omics data. MDV is a web-based interactive analytics and visualisation tool that allows a wide variety of complex datasets to be viewed, annotated, and shared. For example, MDV allows access and integration of data from multi-omic singlecell, spatial imaging, and genomic projects via its user-friendly graphical interface. While MDV streamlines many aspects of data visualisation, tasks that require repetitive actions across multiple features (e.g., constructing charts for visualising gene expression for many genes across a disease condition) may introduce interaction overhead. Such use cases could benefit from batch automation, yet achieving this still typically requires programming knowledge.

Graphical user interfaces (GUIs), although designed for accessibility, can become overcomplicated and inconsistent in functionality. Meanwhile, command-line tools and Application Programming Interfaces (APIs) provide greater flexibility, however they are often highly specific to certain types of data and may alienate non-programmers. This usability gap continues to slow analysis workflows, limit reproducibility, and hinder broader data interpretation.

Recent advances in natural language processing (NLP) and large language models (LLMs) [4, 5, 6] have enabled the development of intuitive, conversational interfaces for interacting with data. Opensource Python frameworks such as LangChain [7] have been developed to facilitate the creation of robust, modular LLM-powered applications. Selected functionalities offered by the LangChain framework encompass agentic systems capable of operating at varying degrees of autonomy [8]. These systems facilitate complex decision-making processes by dynamically invoking external tools, such as a Python Read-Eval-Print Loop (REPL)—an interactive environment for executing code in real time. The usefulness of agentic systems has been demonstrated by their application in tools such as FlowAgent [9] and CellAgent [10], allowing agent-based automation for single-cell RNA-sequencing (scRNA-seq) data analysis. In addition, LangChain supports retrieval-augmented generation (RAG) [11], a paradigm that enhances LLMs outputs by incorporating pertinent information retrieved from external knowledge sources directly into the model’s input prompt. This integration yields responses that are more accurate, contextually grounded, and aligned with the information landscape relevant to the task.

By leveraging these technologies, we have developed **ChatMDV**, a natural language interface for MDV, that enables users to generate, edit, and explore interactive biological visualisations using plain language prompts—without needing to write code. When a natural language prompt is submitted to ChatMDV through the interactive chatbot interface, ChatMDV generates the relevant python code that is executed on the server, generating a new MDV **view** containing one or more **chart**(s) addressing the question. The charts are interlinked and may be cross-queried and filtered. ChatMDV also suggests biological questions that can be queried and visualised using the data.

ChatMDV consists of three modules. Firstly, a custom LangChain [7] agent that understands the type of chart(s) most suited to address the question and the appropriate parameter names from the columns (comprising the dataset) required to build the chart(s). This is passed as a string to the second module comprising ChatMDV, the RAG Pipeline [11] module, dynamically filling the template prompting the module. The RAG Pipeline module, informed by the visualisation type passed on by the custom LangChain [7] agent, is focused on selecting semantically similar Python code templates and passing them to the third and last module. The last module is dedicated to generating the Python code to be executed. The LLMs supporting the framework require no fine-tuning and ChatMDV may be adapted for other dataset modalities (such as spatial transcriptomics or clinical tabular datasets) beyond scRNA-seq data, which is the focus of this study. When the RAG module is called, a vector database managing embeddings of Python code is queried, and based on semantic similarity with the natural language query, the top 5 most similar Python scripts are retrieved, augmenting the prompt with the relevant, retrieved context, informing the LLM with additional details allowing a more focused, accurate, and factually correct response. In addition to the framework powering ChatMDV, we also release a custom Python API which can be used to generate MDV visualisations programmatically, either through the command line or Jupyter notebooks.

To our knowledge, ChatMDV is the only platform that enables users to interactively visualise and further interrogate their -omics (transcriptomics, proteomics, and genomics) data using natural language within a unified analytical interface. While tools such as OMEGA [12] for Napari [2] have begun exploring similar directions in natural language-enabled exploration for image-based biological data, ChatMDV is the first to bring this paradigm to richly annotated single-cell datasets and not only microscopy images, providing a chatbot interface for scalable, domain-aware data visualisation and further analysis.

## Methods

### Design of ChatMDV

ChatMDV is composed of three modules that operate in sequence to interpret user queries and generate interactive charts visualised in MDV the Data and Chart planner agent module, the Retrieval Augmented Generation (RAG) Pipeline module and the Code Generation Chain module (Figure 1). Collectively, these modules implement a combined agentic RAG framework optimised for adaptable, scalable, and reproducible biological data visualisation via structured task decomposition. Complex user queries are broken down into a sequence of manageable subtasks such as query interpretation, data field identification, chart type selection, code synthesis, and visualisation generation which are handled by the specialised modules within the system. Additionally, a memory and history functionality in the Data and Chart Pipeline module supports conversational context and continuity during multi-step graph generation and iterative graph improvement (Figure 1).

**Figure 1.**
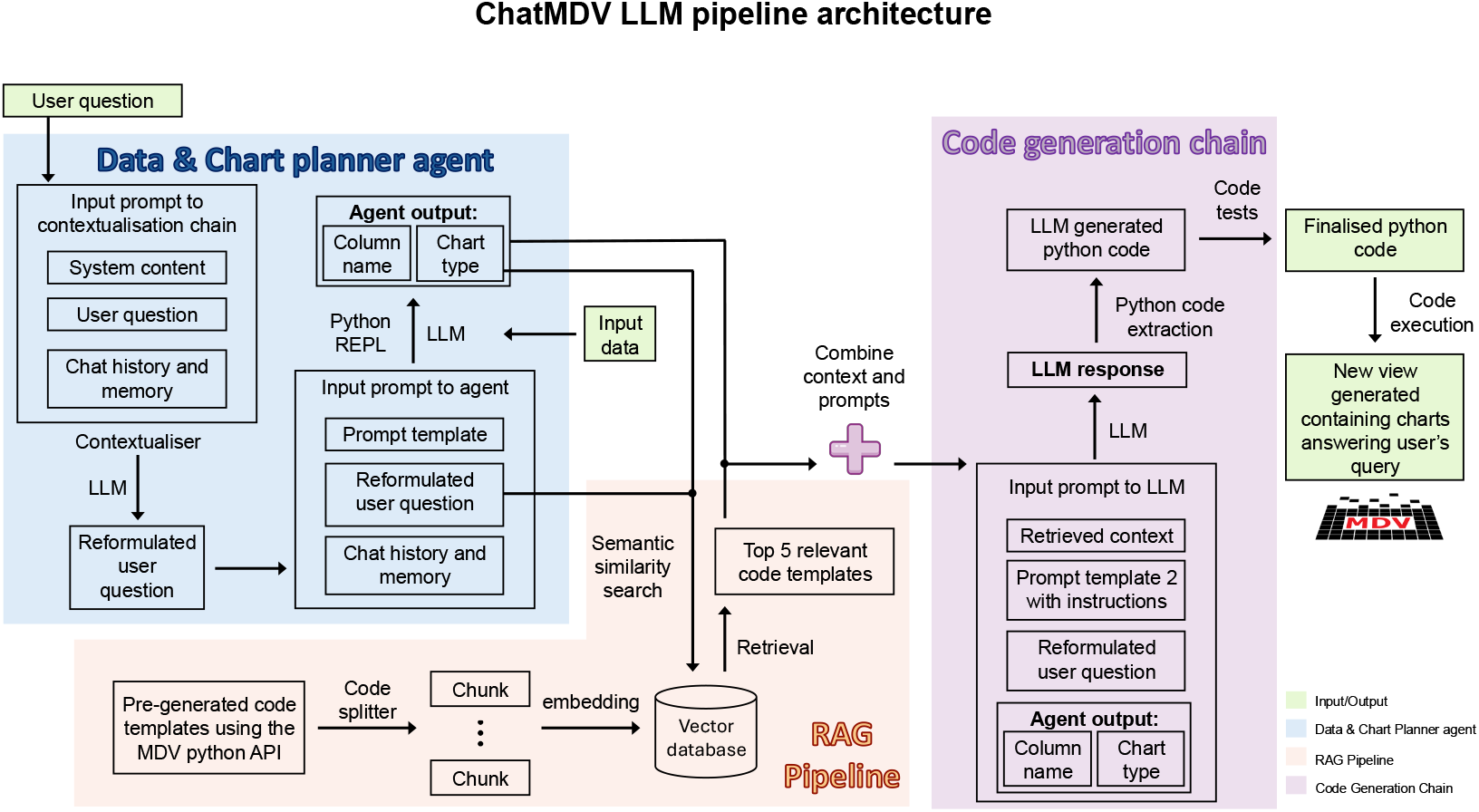
System Architecture of the ChatMDV Pipeline. ChatMDV comprises three core modules: the *Data and Chart planner agent*, the *Retrieval Augmented Generation (RAG) Pipeline* and the *Code Generation Chain*. The *Data and Chart planner agent* module is responsible for interpreting user prompts, identifying the appropriate dataset components and selecting the most suitable chart types for visualisation according to the dataset and question types. This agent also leverages conversational history to contextualise queries based on prior interactions and it is able to autonomously call a Python Read-Eval-Print Loop (REPL) interface to execute code. The suggested chart types and the contextualised user prompt are then forwarded to the *RAG pipeline* module. This module retrieves the five most relevant Python code templates based on similarity from a set of pre-generated code templates generated using the MDV Python API, stored in a vector database. Subsequently, the five most relevant templates, along with the identified dataset components, the suggested chart types and the user’s contextualized query, serve as context input to the *Code Generation Chain* module. This module synthesises and executes the Python script generated. The resulting code and its corresponding view are then returned and rendered within the MDV interface, completing the user query with both executable code and visual output.

MDV and ChatMDV are deployed within an isolated Docker container environment, which mitigates potential security risks by sandboxing execution and preventing access, deletion, or manipulation of the user’s local data.

### Data and Chart Planner Agent

This module initiates the pipeline by interpreting the user’s natural language prompt and by reformulating the user prompt according to conversation history and the system context. Once the reformulated query is submitted to the agent, the LLM-driven agent using its language comprehension capabilities and using the Python REPL interface, interrogates the dataset and identifies the relevant data fields from the input dataset. Additionally, instructed by the prompt framework, it recommends appropriate chart types based on the data fields’ data types. In summary, it performs two primary functions: (i) semantic parsing of the query, and (ii) retrieval of the relevant data fields coupled with chart type suggestions.

The prompt framework underpinning this module was based on the LangChain [7] documentation and the essential prompt elements manually crafted through iterative refinement and enhanced using ChatGPT 4.1. These will be made available in the GitHub repository for this project.

An example prompt (also seen in Figure 2) is “Plot a scatter plot of the UMAP components”. This guides the module to search the dataset for the “UMAP components”. As the agent has access to the dataset, it can search it within the Python REPL environment using the Pandas package [13] and retrieve the exact column names referring to UMAP components. As there is a mention of graph in the prompt, the agent would return the “scatter plot” as a suggestion along the column names. This allows the agent to provide the exact column names and limits hallucinations. If the initial column retrieval is not satisfactory to the agent, the agent retries using an iterative refinement strategy until it returns a satisfactory output or reaches the maximum iteration limit.

**Figure 2.**
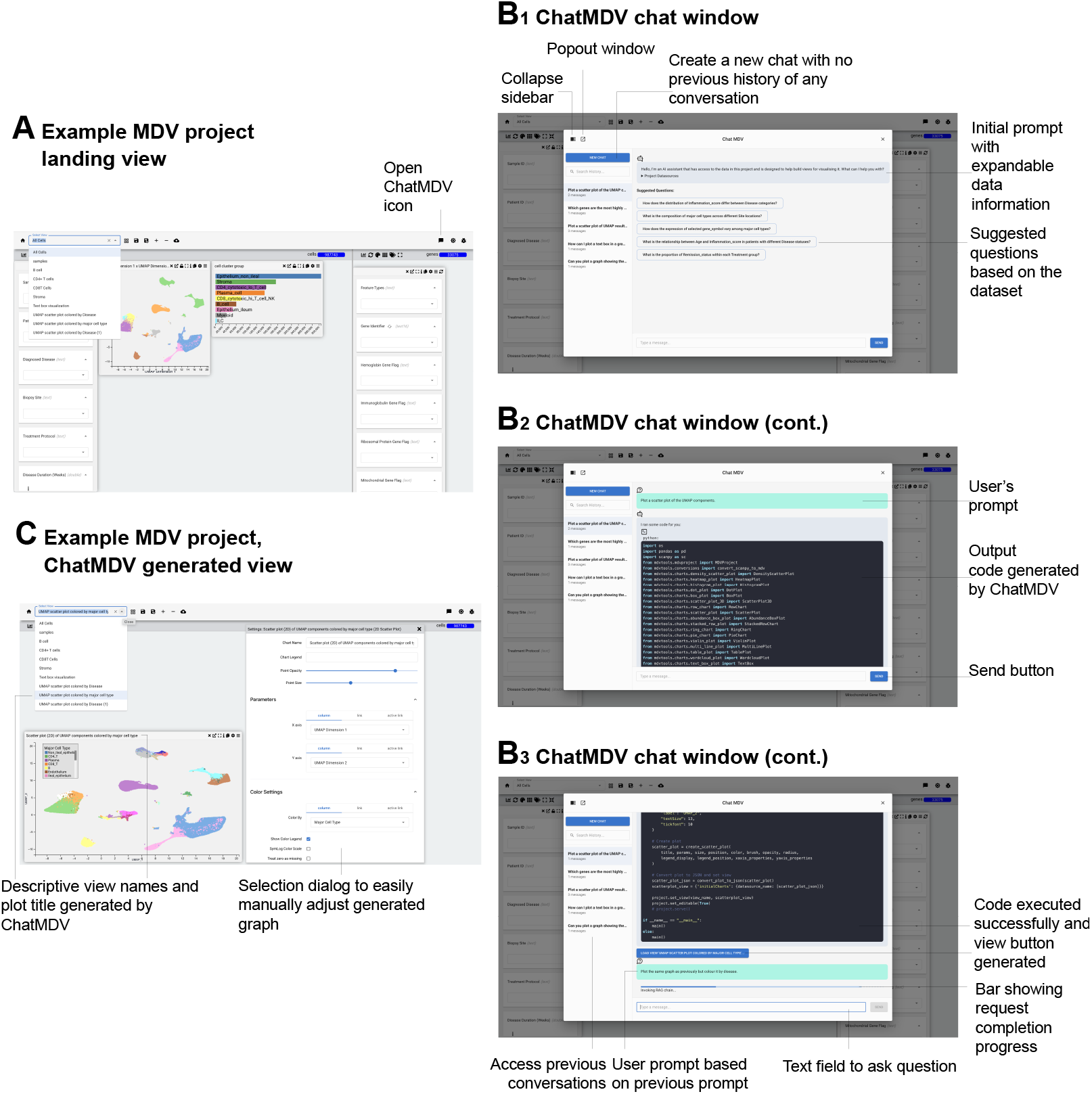
Multi Dimensional Viewer (MDV) user interface (UI) and ChatMDV interaction flow. **A**. Landing page of an MDV project displaying multiple charts, with a dropdown menu listing available views to browse. The ChatMDV icon is shown and it is accessible for initiating an interactive session. **B1**. Upon clicking the ChatMDV icon, a chat window appears with a collapsible sidebar and optional pop-out mode. The initial ChatMDV message appears prompting the users to ask a question. A set of questions to ask ChatMDV is suggested and information about the dataset can be expanded. **B2**. Users can submit natural language queries; ChatMDV outputs the Python code that has been executed. Conversation history is preserved, and a *new chat* button allows starting a fresh session. **B3**. A progress bar indicates execution status of submitted queries, along with real-time feedback on ChatMDV’s reasoning. The *view* button becomes visible after scrolling. Previous conversations can be accessed through the side bar. **C**. The generated visualisations appear upon clicking the *new view* button. Descriptive view names and plot titles aid interpretation, adding an extra layer of information helping the user comprehend the content of the view. The MDV point-and-click interface further supports interactive chart editing enabling even more flexibility for the user.

By generating a targeted list of columns grouped by visualisation intent, this module significantly reduces overhead and irrelevant data handling downstream.

### Retrieval Augmented Generation Pipeline

To populate the vector database in the RAG pipeline module, we developed a library of Python code templates, one per chart type supported by MDV, along with additional templates that combine multiple different chart types. The pre-generated Python code templates are developed using the MDV Python API, an API that enables technically proficient users to programmatically construct, customize, and deploy their MDV projects composed of interactive views created based on their datasets. These templates are data-agnostic, ensuring compatibility across diverse datasets and promoting generalization, reusability, and flexibility. Each code template is segmented into modular code chunks and embedded into vector representations (Figure 1). These embeddings are indexed within an open-source FAISS vector database [14], enabling fast retrieval of relevant code snippets at inference time (Figure 1). The five templates most semantically similar to the reformulated user question and the suggested chart types are retrieved and passed as contextual information to the Code Generation Chain module (Figure 1). Adapting a RAG pipeline in the ChatMDV system architecture supports the next module (Code Generation Chain module) to generate MDV API aligned Python scripts, minimises hallucinations in the LLM responses, provides a flexible mechanism to support new chart types as the MDV architecture evolves and eliminates the need for fine-tuning the LLM, resulting in a streamlined pipeline. Continuing with the example prompt, “Plot a scatter plot of the UMAP components” this query together with the structured output from the previous module — is passed to the RAG pipeline module. In response, the module retrieves the top five most semantically relevant Python code templates from the indexed repository, which in this case includes templates associated with scatter plot visualisations.

### Code Generation Chain

Following the identification of relevant data fields, the chart types suggestion, and the selection of the most semantically similar pregenerated Python code templates, this information is passed as contextual input to the subsequent Code Generation Chain module. This module employs a context-augmented generation strategy to synthesize executable Python visualisation scripts, based on the MDV API. By incorporating the retrieved data fields, recommended chart types, and representative MDV API-based code templates into the prompt, the system is guided to produce code that is both semantically aligned with the user’s intent and syntactically compatible with the MDV framework (Figure 1). For example, for the query “Plot a scatter plot of the UMAP components”, the scatter plot template passed as context will be prioritised by the LLM and the generated code modelled on the template. The UMAP related column names will be used along with customised view names, title names and chart colours.

Using the augmented prompt, the LLM generates a response containing a Python code script based on the pre-generated templates. However, the script contains dataset-specific fields, adjusted axis labels and chart titles, a comprehensive view name, dataset path and addresses the visualisation request put forward by the user. The next step in the module sequence is Python code extraction, as seen in Figure 1, which is performed using a bespoke script run on the server. The script is executed on an isolated Python process within the MDV hosting environment and standard output (successful run) and standard errors are captured. The resulting execution logs are also obtained. The final Python script generates output in JSON format, compatible with MDV rendering.

As shown in Figure 1, the system employs a custom Python execution harness that serializes the target code to a temporary file, executes it in an isolated Python subprocess, and captures standard output and error streams. The resulting execution logs are then programmatically evaluated to verify successful completion within the MDV hosting environment.

This enables visualisations to be served interactively within the MDV interface, supporting direct manipulation of charts and sharing. The Code Generation Chain module is primarily tasked with synthesizing Python scripts aligned with the MDV API, guided by context augmentation derived from relevant data fields, chart types, and pre-generated Python code templates. While the code generation uses the MDV framework, the underlying LLM also has access to widely used Python-based single-cell gene expression analysis libraries such as Scanpy [15]. As a result, when user queries require additional analytical steps—such as normalisation or filtering based on gene expression—the model can incorporate these operations into the generated script while maintaining compatibility with the MDV framework. The choice of Python as the target language further enables integration with a broad ecosystem of scientific libraries, enhancing the analytical flexibility and extensibility of the platform.

Together, these modules allow ChatMDV to convert biological questions into visual answers—automatically, interactively, and reproducibly.

### Multidimensional Viewer User Interface and ChatMDV

ChatMDV is fully integrated into the MDV user interface, enabling users to combine the flexibility of natural language driven view generation with MDV’s interactive, point-and-click visualisation environment. This integration allows users not only to generate initial visualisations using the interactive chat interface but also to further refine or extend them using MDV’s native tools, creating additional views or modifying existing ones for deeper data exploration.

To use ChatMDV, a user clicks on the ChatMDV icon on the top right of the user interface when in a project (Figure 2A). ChatMDV automatically generates a set of suggested questions to provide a starting point for interaction and provide ideas for dataset exploring (Figure 2B_1_). To further help the user understand the dataset, we show an expandable summary of the data columns, types and value ranges (Figure 2B_1_). Once the user asks a question, they can submit the request using the send button (Figure 2B_2_). ChatMDV shows a progress bar along with other useful information regarding the advancement of the query (Figure 2B_3_). Following successful output generation, the Python code is displayed and a view button is generated which directs the user to the new view containing the requested chart(s). The ChatMDV pipeline architecture is designed to maintain awareness of conversation context, enabling a user to ask follow up questions in a conversational matter incorporating conversation history. For example, after generating a view, a user can ask subsequent queries to modify the visualisations, such as asking to add additional genes in a gene expression plot without needing to restate the full request (Figure 2B_3_). Previous conversations can be accessed through a collapsible side bar on the left hand side of the user interface (UI) (Figure 2B_3_). MDV’s rich graphical interface allows users to customize output visualisations through various chart settings, including point size, colour scheme, axis dimensions, and legend position. (Figure 2C).

### Instance and project creation

MDV instances can be locally installed by cloning our GitHub repository https://github.com/Taylor-CCB-Group/MDV/tree/dev and setting up the appropriate Docker containers. ChatMDV can be enabled by providing an OpenAI key. Guidance and instructions can be found in our website https://mdv.ndm.ox.ac.uk/ and our GitHub repository: https://github.com/Taylor-CCB-Group/MDV/tree/dev. Once a Docker container is set up, a user can upload their scRNA-seq.h5ad object, an example dataset is provided in our website along with more detailed instructions on uploading your own dataset.

Once the scRNA-seq dataset or project is uploaded, the user can access both the cells’ dimension and the genes’ dimension stored in Anndata objects [16], a data structure used in single-cell analysis which stores the gene expression matrix along with cell and gene level metadata. Linking the gene expression matrix with the associated cell and gene metadata is performed automatically as part of the MDV project creation. This complexity is abstracted from the user and this automation reduces the need of manual data wrangling reducing the computational overheard for the users. Therefore, a user can access the data and generate more complex, biologically relevant and informative charts. Gene identifiers such as gene IDs or gene names are used to link the data.

## Evaluation and Case Study

### Datasets for ChatMDV Evaluation

To demonstrate the usability and practical utility of ChatMDV in data exploration and visualisation, we selected single-cell RNA-sequencing (scRNA-seq) [17] as a representative modality. scRNA-seq datasets are characterized by their high dimensionality, cellular diversity, and increasing ubiquity in modern biomedical studies, making them an ideal benchmark for assessing the capabilities of ChatMDV. To this end, we selected three publicly available scRNA-seq datasets of varying biological and technical complexity, which served as the basis for constructing domain-relevant queries to evaluate ChatMDV’s functionality.

The first dataset is the widely used Peripheral Blood Mononuclear Cells 3K dataset (PBMC3K) of 10X Genomics [18]. It includes transcriptomic profiles from 2,700 immune cells isolated from a healthy donor. This benchmark dataset is widely used for cell clustering, dimensionality reduction, and gene expression profiling. Its manageable size and well-characterised annotations make it an ideal testbed for validating ChatMDV’s core visualisation capabilities, including scatter plots of UMAP projections, gene expression heatmaps and violin plots. The second dataset is a comprehensive lung cancer atlas from the Human Cell Atlas [19], offering a rich landscape of cell states across healthy and diseased lung tissue, at a single time point. It includes 2,282,447 cells from the respiratory tract of 486 individuals [19]. The third dataset is the TAURUS dataset [20] which comprises longitudinal scRNA-seq profiles from patients with Crohn’s disease (CD) and ulcerative colitis (UC) undergoing anti tumor necrosis factor (anti-TNF) therapy. It captures samples from pre- and post-treatment timepoints and includes subclustered cell states annotated across multiple disease and response groups, for 987,743 cells.

Each dataset introduces an additional layer of complexity. We begin with the PBMC3K [18, 21] dataset, a relatively small and well-characterised reference dataset. The lung cancer atlas dataset [19] adds complexity through its cross-sectional design, capturing cells from multiple individuals and tissue/cell types. The TAURUS [20] dataset adds a longitudinal component, consisting of samples collected from multiple individuals at multiple timepoints, both before and after anti-TNF treatment. This enables the construction of more advanced, biologically contextualised queries to prompt ChatMDV for each dataset.

### Evaluation Approach

To validate ChatMDV’s capabilities, we developed a systematic approach for testing and assessing the system’s performance. It comprised a set of domain specific questions to simulate typical exploratory biological analysis with the goal of elucidating data properties and aid in scientific discovery and hypothesis generation. Natural language prompts were written to resemble domain-specific questions a typical bioinformatician or experimentalist might pose, such as comparing gene expression across cell states or visualising treatment effects over time or for different patients. As part of the assessment, an automated evaluation system was built to submit the set of user queries to ChatMDV and generate responses addressing each query as distinct views within a new MDV project for each dataset (Figure 3A). The generated views were then manually reviewed and annotated, and assigned ratings on a 5 point scale, ranging from 1 (View not generated or no charts present) to 5 (Perfect view). These ratings are summarised in Figure 3B and are described in detail in Section Output Classification and Evaluation Criteria. Additionally, each question was assigned a complexity score (ranging from 1 to 7) reflecting its semantic and analytical difficulty, which was found to impact ChatMDV’s performance. More details on the complexity scoring scheme are provided in Section Prompt Design and Complexity Categorisation and also seen in Figure 3C.

**Figure 3.**
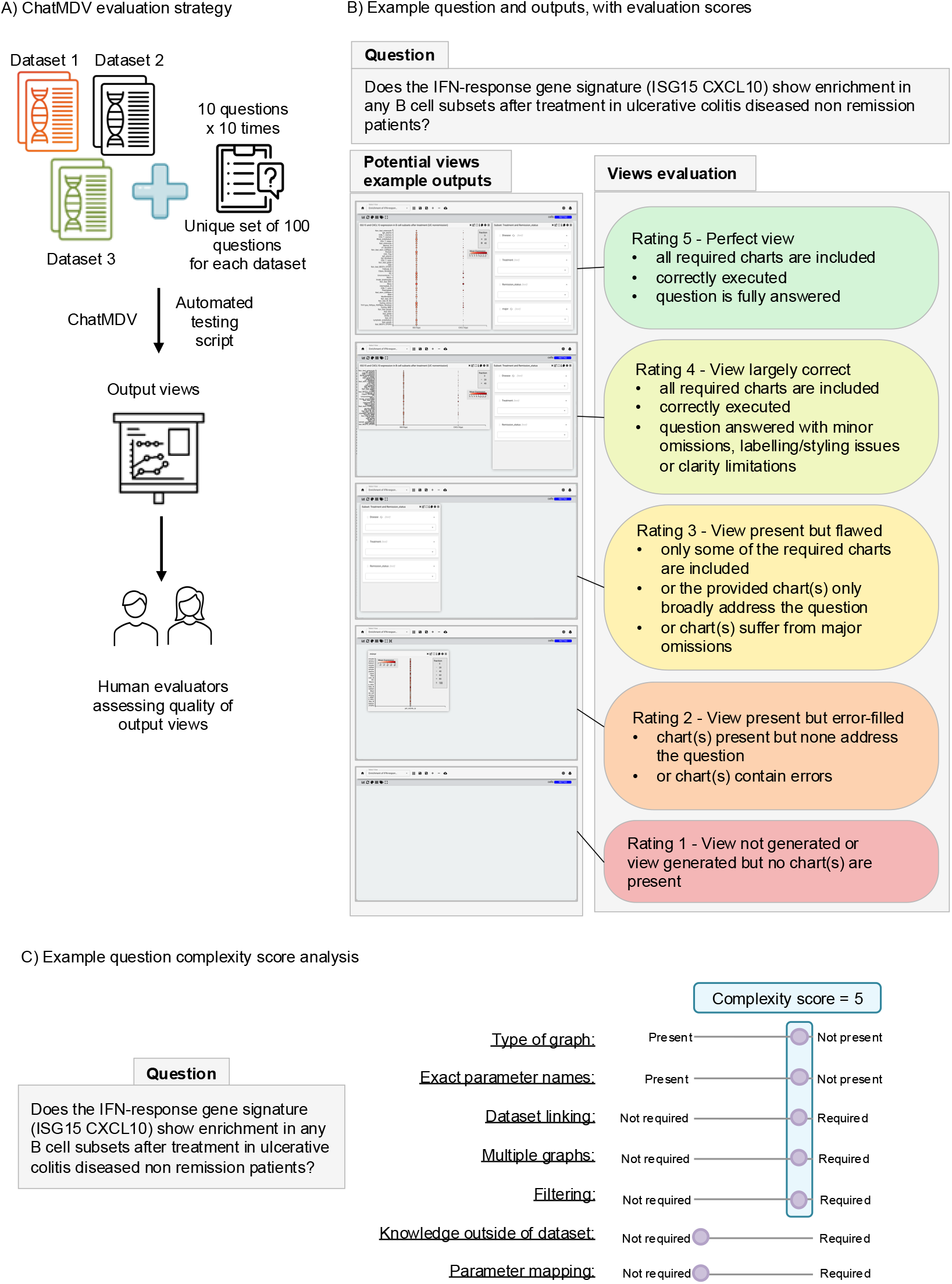
ChatMDV evaluation strategy, evaluation example and complexity score assignment example. **A**. ChatMDV was evaluated using three distinct scRNA-seq datasets to demonstrate its generalisability and broad applicability. For each dataset, a set of 10 unique natural language questions was curated and each question was submitted 10 times using an automated Python evaluation testing script. The resulting visualisations were generated within the same MDV project, assessed and scored. **B**. Example of a question submitted to ChatMDV, with representative visualisation outputs and corresponding qualitative evaluation ratings. **C**. Complexity analysis and complexity score assignment example for the example question presented in panel B.

### Prompt Design and Complexity Categorisation

Multiple test prompts were developed for each data set to ensure coverage of a broad range of biological use cases. The prompts (which are reported in Figure 4 and in Supplementary File 1) were developed through a combination of expert consultations, a detailed review of the relevant scientific literature accompanying each dataset and ChatGPT’s assistance. During the creation process we made sure that the prompts were biologically relevant, spanned a range of graphs supported by MDV and varied in complexity. We assessed the complexity of each test prompt by using a bespoke complexity schema. We assigned a complexity score to each test prompt by assigning a point to the overall score based on whether the prompt meets the following criteria:

**Figure 4.**
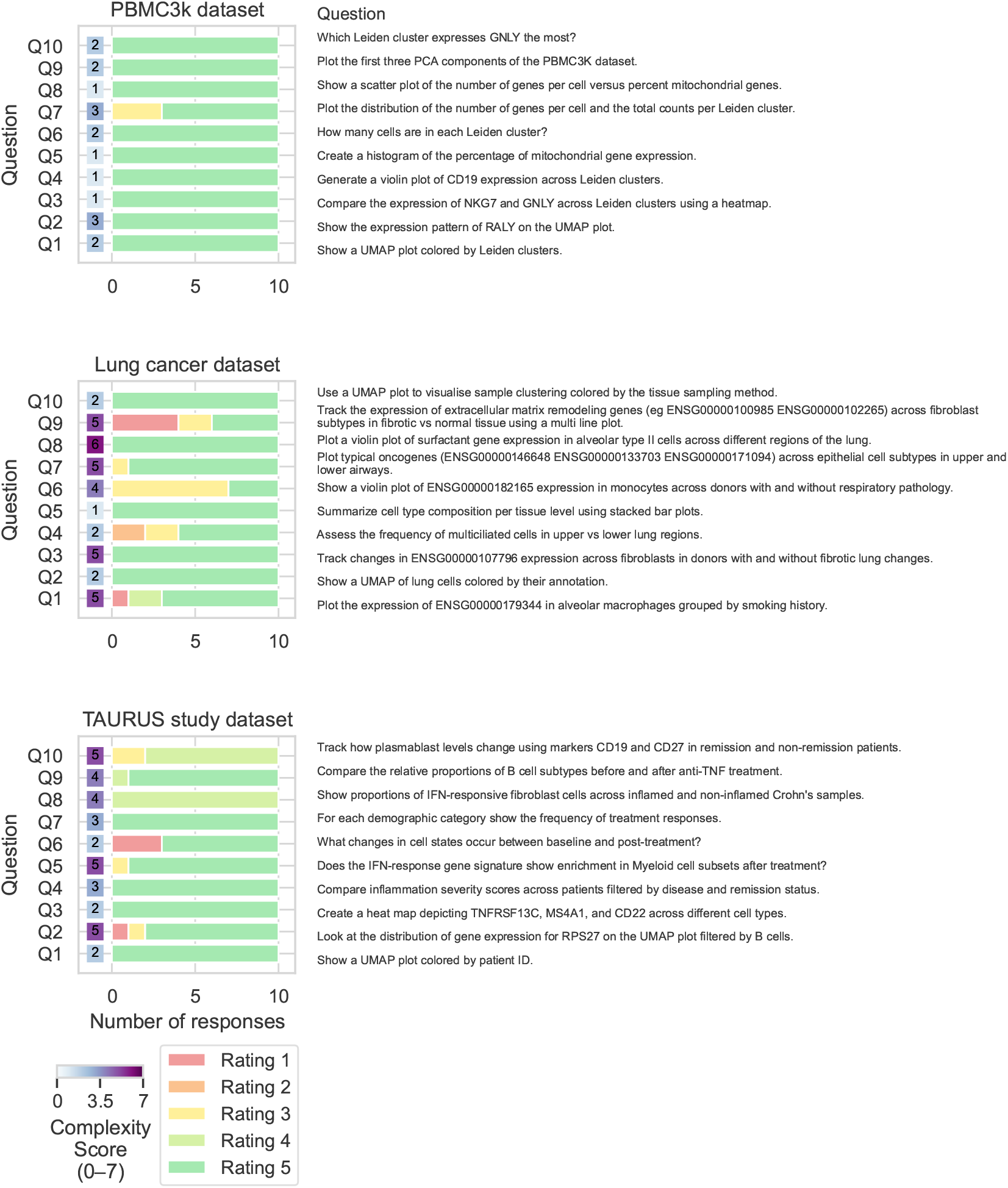
ChatMDV evaluation results. Summary of evaluation outcomes presented as bar plots, shown for each dataset. Evaluation questions are shown in the right hand side of the y-axis with corresponding complexity score on the left hand side. Each question is run through the automated evaluation script 10 times, and the outcomes are annotated using a rating scale, ranging from “Rating 5” (which corresponds to a “Perfect view”) to “Rating 1” (corresponding to an empty view or a view that is not generated). Next to the evaluation outcomes barplots the complexity score is shown, with the most complex question being assigned a value of 7 and the least complex is assigned a value of 1. Detailed complexity scoring criteria for each question and complexity assignment are provided in Figure 3 and Supplementary File 1. The breakdown of the annotation results per dataset is provided in Supplementary File 3.

- **Type of graph:** The prompt explicitly states the desired graph type (e.g., bar chart, line plot, scatter plot). Omitting this information increases the complexity score by one point.
- **Exact parameter names:** The prompt includes the precise column or gene names as they appear in the dataset, for all the required data fields. If exact field names are not provided, a complexity point is added.
- **Dataset linking:** If the prompt requires combining information from different data sources (e.g., linking gene expression data with metadata using gene IDs), one point is added.
- **Multiple graphs:** The prompt expects the generation of more than one graph, which adds an additional complexity point.
- **Data filtering:** If the prompt includes conditions that require filtering or selecting specific subsets of the data (e.g., only patients with a certain disease), one point is added.
- **Knowledge outside of the dataset:** If the prompt refers to information not directly available in the dataset (e.g., referencing a class of genes without listing them), it adds complexity.
- **Parameter mapping:** The LLM has to identify that the information in the prompt requires mapping to a different representation from the dataset. This additional reasoning step is adding to the prompt’s complexity. For example, if a gene is referenced by name in the prompt but is stored as an Ensembl ID in the dataset [22], we assign an additional complexity point.

After the set of 10 questions was constructed for each dataset, each question was repeated 10 times resulting in a set of 100 natural language prompts to be tested for each use case (Figure 3). The complete evaluation pipeline was executed across all prompts. For each prompt, ChatMDV’s output was documented, including the output of the “Data and Chart Planner Agent”, the top 5 relevant code templates retrieved and passed as context, and the output code generated and executed by ChatMDV for each natural language prompt. Potential execution errors were also recorded for further understanding, debugging and improving the pipeline. The full output of ChatMDV for each natural language prompt is given in Supplementary File 2.

This fully automated evaluation pipeline allowed us to systematically assess ChatMDV’s capacity to understand varying query structures and intent. It helped us assess reproducibility of results and robustness of the system. Moreover, it helped improve the pipeline by prompt engineering and experimenting with the both the agent and RAG systems’ parameters. The iterative improvement process was performed with different prompts which spanned from very low complexity prompts such as “Plot a scatter plot of the number of genes versus the number of cells” and reached high complexity prompts such as “Plot the gene expression for the highest expressing genes in the dataset”.

### Output Classification and Evaluation Criteria

The output of the evaluation pipeline was evaluated using a pre designed rating system to assess the completeness, quality, and relevance of each generated view (Figure 3). A rating of 5, ‘Perfect view’, was assigned to the view when all required charts were present, the code generating them correctly executed, and fully answered the natural language prompt. A rating of 4, ‘View largely correct’, indicated that all necessary charts were included and accurate, but with minor omissions, clarity issues, or styling problems. A rating of 3, ‘View present but flawed’, reflected partial success, for example, only some required charts were present, or the charts only broadly addressed the question, or suffered from major omissions. A rating of 2, ‘View present but error-filled’, was given when charts were present but did not address the question or contained significant errors. Finally, a rating of 1 indicated that no view was generated or an empty view was generated without any visual content. Common reasons for this included: failure to generate Python code (i.e., the pipeline did not complete), Python code generation followed by execution errors, or successful execution of Python code that failed to produce valid graphs due to runtime errors. Summarising the results, error classifying and investigating the reasons behind failed chart/view generation enabled targeted system improvements and informed development priorities. Gradual improvements included prompt engineering, optimising conditions related to the RAG pipeline such as the similarity threshold, embedding model choice, model temperature and improved error handling and recovery mechanisms. The underlying reasons for not achieving a perfect rating (Rating of 5) included incorrect identification of variable names corresponding to the desired parameters, mismatches between the data type required by the chart type used, failure to recognise and map relevant genes present in the dataset, and omission of selection dialog plots necessary for filtering the data.

## Results

A user can instruct ChatMDV to “plot a chart of the uniform manifold approximation and projection (UMAP) components” of a dataset and ChatMDV can choose a suitable graph (scatter plot) and visualise the user’s request (Supplementary video 1). Moreover, ChatMDV is able to identify the column names from within the dataset that correspond to the “UMAP components” so it can create a visualisation successfully (Supplementary video 1). Users can ask follow-up questions to modify the visualisation (Supplementary video 2), utilising ChatMDV’s capability of retaining conversational history. ChatMDV can also create more than one visualisations within the same view, by asking it “Plot the gene expression of these genes *TP53* (ENSG00000141510), *KRAS* (ENSG00000133703), and *ALK* (ENSG00000171094) in separate box plots for different lung conditions.” (Supplementary video 3). This functionality is particularly useful for automating repetitive graph generation tasks such as creating the same plot type for multiple marker genes. The generated charts can then be interactively adjusted and customised by the user within the MDV interface (Supplementary video 3). The datasets within the project can be filtered and sub setted, for example by asking ChatMDV to “Compare inflammation severity scores across patients filtered by disease and remission status”. ChatMDV adds a selection dialog plot which is MDV’s chart type for filtering the dataset (Supplementary video 4). Adding a text box is a chart functionality supported by MDV. ChatMDV is able to utilise this type of chart to output information about the dataset (Supplementary videos 5 and 6) and beyond. Example natural language prompts include, “Print your understanding about the dataset in a text box.” and “Print summary statistics and other important info about the dataset in a text box.” (Supplementary videos 5 and 6 respectively). ScRNA-seq data is stored in Anndata objects [16], which organise information into multiple matrices, usually separating cell metadata and gene annotations into distinct structured components. Plotting charts showing gene expression related properties require integrating the cells’ and genes’ matrices by linking on gene identifiers. ChatMDV’s capabilities involve linking different matrices on predefined gene identifiers (Supplementary video 7). The linking gene identifier is predefined in the Anndata object and it can be one of gene name or gene ID. ChatMDV can accurately map between gene names and gene IDs, ensuring that the appropriate data is retrieved and the correct plots are generated (Supplementary video 8). The lung cancer dataset uses Ensembl gene IDs [22] as the gene identifier and in Supplementary video 8 the natural language prompt (“Plot the expression of HLA-DQB1 in alveolar macrophages grouped by smoking history”) is querying ChatMDV using the gene name.

By implementing the ChatMDV pipeline with support for executing Python code, we enable the underlying LLM to apply its external domain knowledge and make use of established packages for scRNA-seq analysis such as Scanpy [15], a widely used package in the field of single-cell analysis. In Supplementary video 9, ChatMDV demonstrates this capability by invoking the rank_genes_groups() function to perform differential gene expression analysis, filter statistically significant genes, and identify top candidates based on effect size. It then visualises the expression patterns of the top three genes across cell clusters using both a dot plot and a heatmap plot. The example demonstrated in Supplementary video 9 uses the PBMC3K dataset, whose.h5ad file is approximately 32 MB in size. In comparison, the TAURUS and lung cancer datasets are three orders of magnitude larger. Running the described scRNA-seq analysis on datasets exceeding 4 GB requires hardware with significantly more random-access memory (RAM), than the ones we use to support the projects presented in this publication. For this reason, the analysis is demonstrated on the smaller PBMC3K dataset.

To demonstrate ChatMDV’s robustness and evaluate its performance, we used the automated evaluation pipeline described in Sections Prompt Design and Complexity Categorisation and Output Classification and Evaluation Criteria. A summary of results is presented in Figure 4 with a detailed breakdown available in Supplementary File 3. The sets of 100 prompts for each dataset are independent test sets, not used in iterative refinement. We achieved a success rate (any rating except rating 1) of 100% for the PBMC3K dataset, 95% for the lung cancer dataset and 96% for the TAURUS dataset. The success rate is defined as the ratio of successful attempts versus all ten attempts per question averaged for the questions per dataset.

We observe that all natural language prompts receiving a rating of 1 had a complexity score greater or equal to 2. This suggests that ChatMDV achieves a 100% success rate on simpler prompts, specifically those that clearly describe the desired chart type and explicitly state the relevant parameter names. We rank the datasets by increasing biological complexity as PBMC3K, lung and TAURUS, as TAURUS is a longitudinal scRNA-seq dataset.

The results indicate that ChatMDV’s performance, measured by the success rate, is slightly reduced for the lung cancer dataset. We hypothesize that this is due to a combination of factors, including increased dataset size (PBMC3K < 100 MB; lung cancer ∼22 GB; TAURUS ∼12 GB) and the number of columns available to the ChatMDV pipeline (PBMC3K: 12 columns, lung cancer: 76 columns, TAURUS: 42 columns). Additionally, we note that the TAURUS dataset features more comprehensively curated column names, with clear and descriptive labels, whereas the lung cancer dataset lacks consistent column name curation. The higher number of columns in the lung cancer dataset also increases the search space for the Data and Chart Planner Agent, making it more prone to errors and increasing execution time.

## Discussion

ChatMDV introduces a novel approach to biological data visualisation using natural language. Building on MDV’s established visualisation functionality it provides a compelling way to explore large and complex single-cell data sets. Through the use of large language models (LLMs) and a sequential retrieval-augmented generation (RAG) pipeline, ChatMDV provides suggested questions driven by the data to provide a starting point for analysis. Biological questions such as “Which Leiden cluster expresses GNLY the most?” and “Run the scanpy rank_genes_groups_df function on the data and find the most highly differentially expressed genes.” can be used allowing users to focus on intent rather than learning complex user interfaces or coding. Care has been taken to support reproducibility by ensuring generated code is accessible from ChatMDV’s chat history.

MDV’s user interface allows post refinement of the chart properties (layout, font size etc.), provides a seamless workflow to query data and produce visualisations that can be used in presentations and publications. By coupling LLM driven automation with hands on point-and-click refinement, ChatMDV bridges the gap between code-centric bioinformatics and intuitive tooling making bioinformatics analysis exploration accessible.

The ChatMDV architecture allows a rapid way to add new analysis methods by adding documents or code to the database when needed. Our RAG approach also enhances query transparency by linking generated content back to the source code. This traceability improves user confidence and interpretability which is an important factor for adoption in scientific workflows.

A further strength of ChatMDV lies in its modular design. Because the tool operates by interfacing with Python APIs, it can be adapted to other domains or frameworks beyond MDV with minimal modification. Unlike static visualisation libraries such as Matplotlib or Seaborn, the MDV interface supports interactive, point-and-click editing of plots, streamlining iterative workflows and improving user experience.

Overall, ChatMDV demonstrates how LLM-powered interfaces can democratise access to computational tools and enhance biological data exploration in a transparent, reproducible, and interactive way.

### Future Directions

Tools like FlowAgent [9] show how multi-agent planners can automate scRNA-seq pipelines and general bioinformatics workflows. ChatMDV slots into this ecosystem as the interactive visual layer, which in future could enable a seamless transition from agent generated results to user-driven visual exploration. In this fast moving field, we predict LLM services around biological data types will rapidly increase. Frameworks such as Model Context Protocol (MCP) [23] offer a public, language-agnostic specification that any agent can plug into, improving discoverability and cross-tool portability. Such services as SCMCP hub [24] and MCPmed [25] can be incorporated into MDV’s framework to allow a range of diverse services to be visualised based on natural language.

We see ChatMDV’s RAG architecture facilitating ways of integrating other unstructured biological assets including PDFs, Electronic Lab Notebooks, Jupyter notebook snippets etc. allowing the system to suggest domain aware queries building on the automated suggested queries already available.

Finally open-source LLMs, such as LLaMA [5], source models can be used for local deployment, rather than using commercial artificial intellegince (AI) APIs, prioritising data privacy for groups or organisations that wish to use ChatMDV to interact with sensitive patient data.

### Limitations

ChatMDV occasionally returns syntactically invalid or unparsable Python code. Errors frequently occur when the system misuses data types, such as using a categorical variable as a chart parameter in a chart that requires a numerical one, like scatter or line charts. Other examples include when ChatMDV hallucinates a parameter name or tries to use a gene that does not exist in the dataset. To address this, we implemented extensive error handling, iterative retries, and comprehensive logging to help identify and resolve issues quickly.

When ChatMDV is required to perform more intense scRNA-seq analysis using scanpy, the computer’s RAM in the docker container may be an issue, For example, running a query “Perform differential analysis using the scanpy dataset and find the least expressed genes.” on the lung cancer data set (∼22 GB) in a typical docker container on a desktop (typically 8-12GB) would fail.

## Additional Files

**Supplementary File 1**. Contains the full set of natural language prompts used to evaluate ChatMDV across all three datasets, along with the corresponding complexity scores. Additionally, contains a table indicating which complexity criteria are fulfilled for each prompt, with crosses denoting satisfied conditions.

**Supplementary File 2**. Contains the ChatMDV output per prompt for each dataset shown in Figure 4. The file includes the output of the “Data and Chart Planner Agent”, the top 5 relevant code templates retrieved and passed as contexts, and the output code generated by ChatMDV for each natural language prompt. In cases where a view failed to be generated, corresponding code execution errors are also reported.

**Supplementary File 3**. Contains the detailed breakdown of the evaluation results for each dataset summarised in Figure 4. Annotated rating is given for each prompt run through the evaluation system.

**Supplementary Video 1**. By interpreting the user’s intent, ChatMDV selects appropriate chart types and parameter names to generate a suitable graph. The video has been sped up by a factor of 2.

**Supplementary Video 2**. ChatMDV allows users to interactively modify the generated visualisations through the settings dropdown of the point-and-click interface. The video has been sped up by a factor of 2.

**Supplementary Video 3**. A single natural language query can prompt ChatMDV to produce several charts within the same view. Once generated, the user can further refine and modify the individual charts interactively within the MDV interface. The video has been sped up by a factor of 2.

**Supplementary Video 4**. ChatMDV enables interactive dataset filtering by generating selection dialog plots in response to natural language prompts. The video has been sped up by a factor of 2.

**Supplementary Video 5**. ChatMDV uses MDV’s text box chart functionality to produce textual summaries describing the dataset. The video has been sped up by a factor of 2.

**Supplementary Video 6**. ChatMDV generates summaries of dataset statistics using MDV’s text box chart type. The video has been sped up by a factor of 2.

**Supplementary Video 7**. ChatMDV is able to link data from distinct matrices stored in the Anndata object. In this example, the linking of gene expression is demonstrated by colouring the scatter plot showing the UMAP components by the expression of the *RPS27* gene for the TAURUS dataset. The video has been sped up by a factor of 2.

**Supplementary Video 8**. Not all linking identifiers are gene names, but they can be gene IDs too. In this example, ChatMDV automatically determines the correct linking identifier, gene ID in this case, even though the user’s prompt refers to the gene by name, demonstrating its ability to map between different linking fields. The video has been sped up by a factor of 2.

**Supplementary Video 9**. ChatMDV performs differential gene expression analysis using rank_genes_groups() from Scanpy [15], filters significant genes by effect size, and visualises the top three across cell clusters using a dot plot and heatmap. The video has been sped up by a factor of 2.

## Abbreviations

AI: artificial intelligence;
anti-TNF: anti tumor necrosis factor;
APIs: application programming interfaces;
CD: Crohn’s disease;
FAIR: Findability Accessibility Interoperability and Reuse;
GUIs: graphical user interfaces;
LLMs: large language models;
MCP: model context protocol;
MDV: Multi-Dimensional Viewer;
NLPs: natural language processing;
PBMC3K: peripheral blood mononuclear cells 3K;
RAG: retrieval augmented generation;
RAM: random-access memory;
REPL: Read-Eval-Print Loop;
scRNA-seq: single-cell RNA sequencing;
UC: ulcerative colitis;
UI: user interface;
UMAP: Uniform Manifold Approximation and Projection;

## Author Contributions

Maria Kiourlappou: Visualisation, Project Administration, Investigation, Formal Analysis, Conceptualization, Methodology, Validation, Writing - Original Draft Preparation, Writing - Review & Editing, Data Curation. Peter Todd: Software, Methodology, Validation. Yaxuan Kong: Investigation, Methodology, Writing - Review & Editing. Jayesh Hire: Software. Furquan Nawab Mohammed: Software. Martin Sergeant: Software, Data Curation. Stefan Zohren: Supervision, Writing - Review & Editing. Jim Hughes: Supervision. Stephen Taylor: Funding Acquisition, Supervision, Project Administration, Conceptualization, Validation, Writing - Review & Editing, Resources

## Acknowledgments

We are grateful to Christopher Buckley, Calliope A. Dendrou, and Devika Agarwal, the collectors and curators of the TAURUS study dataset for their valuable help, guidance, and insights provided regarding the dataset. We also thank Mostafa Ibrahim for discussions and feedback on early stages of the project. We also acknowledge the assistance of ChatGPT in refining the manuscript language and the code.

## Availability of Source Code and Requirements

- **Project name:** ChatMDV
- **Project page:** https://github.com/Taylor-CCB-Group/MDV/tree/dev
- **Operating system(s):** Platform independent (tested on Linux and macOS)
- **Programming language:** Python / Javascript / Typescript
- **Other requirements:** OpenAI API key, MDV (latest release), Docker (for local install)
- **License:** MIT License
- **Containerisation:** A Docker container supporting the full pipeline is available in our GitHub repository https://github.com/Taylor-CCB-Group/MDV/tree/dev
- **Restrictions to use by non-academics:** None

